# THC modifies the impact of heroin delivered by vapor inhalation in rats

**DOI:** 10.1101/2021.04.26.441541

**Authors:** Arnold Gutierrez, Jacques D. Nguyen, Kevin M. Creehan, Mehrak Javadi-Paydar, Yanabel Grant, Michael A. Taffe

## Abstract

Opioids are effective medications, but they have several key limitations including the development of tolerance, establishment of dependence, diversion for non-medical use and the development of addiction. Therefore, any drugs which act in an additive or synergistic fashion with opioids to address medical applications have the potential to reduce opioid-related harms. This study was conducted to determine if heroin and Δ^9^-tetrahydrocannabinol (THC) interact in an additive or independent manner to alter nociception, body temperature and spontaneous locomotor activity when inhaled or injected.

Groups of male and female rats implanted with radiotelemetry transmitters were exposed to vapor for assessment of effects on temperature and activity. Heroin (50 mg/mL in the propylene glycol; PG) inhalation increased temperature and activity whereas THC (50 mg/mL) inhalation decreased temperature and activity. Effects of combined inhalation were in opposition, and additional experiments found the same outcome for the injection of heroin (0.5 mg/kg, s.c.) and THC (10 mg/kg, i.p.) alone and in combination. In contrast, the co-administration of Heroin and THC by either inhalation or injection produced additive effects on thermal nociception assessed with a warm water tail-withdrawal assay in male and female Sprague-Dawley and Wistar rats.

The conclusion of this study is that additive effects of THC with heroin on a medical endpoint such as analgesia may not generalize to other behavioral or physiological effects, which may be a positive outcome for unwanted side effects.

## 1. Introduction

Expansion of the use of cannabis for purported medical benefits, including for pain, stimulates interest in the possible opioid-sparing effects of cannabis constituents, including the primary psychoactive compound Δ^9^-tetrahydrocannabinol (THC). While opioids are effective medications, they come with many limitations including the development of tolerance and the need to increase doses for similar efficacy, development of dependence which counter-indicates rapid discontinuation, diversion for non-medical use and the development of addiction. Any drugs which may act in an additive or synergistic fashion with opioids therefore have the potential to reduce opioid-related concerns. There are known interactions by which endogenous cannabinoid receptor 1 (CB_1_) ligands may enhance signaling of mu opioid receptors (Parsons and Hurd, 2015), thus generating a reasonable mechanistic hypothesis for evaluating opioid-sparing effects of cannabis constituents.

Cannabis co-use triples the risk of dependence on heroin in those diagnosed with a substance use disorder (Crummy et al., 2020) and cannabis users over 50 years of age exhibit a 6.3 increased odds ratio of heroin use (Ramadan et al., 2020). Adolescents in one urban setting who use cannabis regularly are at twice the risk for opioid misuse compared with occasional users of cannabis (Reboussin et al., 2020) and age 14 onset of frequent cannabis use increases the risk of opioid use at age 19 (Thrul et al., 2020). Almost two thirds of individuals in one sample first used heroin while co-using cannabis (Olthuis et al., 2013) and 50-60% of individuals in heroin-maintenance and methadone-maintenance treatment for opioid use disorder were co-using cannabis (Musshoff et al., 2010). On a day by day basis, cannabis use doubles the risk of non-medical opioid use in adults with problematic substance use (Gorfinkel et al., 2020). Thus, although cannabis may have the potential to reduce opioid use in medical patients, there is also a clear risk for cannabis increasing the non-medical use of heroin from adolescence into middle age. Consideration of these phenomenon spurs interest in determination of the interactive effects of cannabinoids and opioids across multiple behavioral and physiological endpoints to lend greater context for “opioid sparing” recommendations for cannabis use.

We recently presented evidence that THC enhances the effects of oxycodone in an anti-nociception assay in rats and also the effects of a unit dose of oxycodone or heroin when self-administered (Nguyen et al., 2019). Maguire and France have shown that the nature of the anti-nociceptive interaction (supra-additive, sub-additive, additive) between cannabinoids and opioids may depend on the specific drugs that are involved (Gerak et al., 2019; Maguire and France, 2018; Maguire et al., 2013) which cautions against making generalizations across either class of substances, before data are available. In the case of both nociception and drug self-administration, the effects of opioids and cannabinoids are often in the same direction, i.e., anti-nociceptive (Li et al., 2008; Lichtman and Martin, 1990; Manning et al., 2001; Peckham and Traynor, 2006; Wakley and Craft, 2011) and rewarding (Blakesley et al., 1972; Justinova et al., 2003; Killian et al., 1978; Panlilio et al., 2010; Vendruscolo et al., 2018). This can make it difficult to determine if the outcome of co-administration is due to the additive effects of independent mechanisms or the interaction of signaling within the same mechanistic pathways (Ahmad et al., 2013).

Determination of any heroin / THC interactions for *in vivo* endpoints that are expected to change in *opposite* directions after each drug is administered individually can help to further parse the specificity of any apparent additive effects. We’ve identified conditions under which either inhaled or injected heroin can *increase* the body temperature and spontaneous locomotor activity (Gutierrez et al., 2020a), and conditions under which THC can *decrease* body temperature and locomotor activity (Javadi-Paydar et al., 2018; Nguyen et al., 2016b; Taffe et al., 2020). We’ve further shown that the locomotor effects of nicotine and THC on activity can oppose each other when co-administered (Javadi-Paydar et al., 2019b); effects of each drug to decrease body temperature were also dissociated based on the timing, post-administration. This study was conducted to determine if heroin and THC interact in an additive or independent manner to alter thermal nociception, body temperature or spontaneous locomotor activity when inhaled or injected.

The recent broad availability of e-cigarette style Electronic Drug Delivery Systems (EDDS) supports the possibility of delivering a range of drugs other than nicotine, including opioids, via vapor inhalation. Use of these devices for ingestion of cannabis extracts has become increasingly popular (Allem et al., 2019; Dugas et al., 2020; Kowitt et al., 2019; Nicksic et al., 2020; Pearson and Villanti, 2020). The EDDS can be used to deliver active doses of a wide range of drugs including amphetamine and cathinone derivative psychomotor stimulants (Nguyen et al., 2016a; Nguyen et al., 2017), opioids (Gutierrez et al., 2020a; Moussawi et al., 2020; Nguyen et al., 2019; Vendruscolo et al., 2018), cannabinoids (Breit et al., 2020; Freels et al., 2020; Javadi-Paydar et al., 2019a; Javadi-Paydar et al., 2018; Moore et al., 2020a; Nguyen et al., 2016b) and nicotine (Cooper et al., 2020; Frie et al., 2020; Javadi-Paydar et al., 2019b; Montanari et al., 2020; Ponzoni et al., 2015) to rats and mice; for review see (Miliano et al., 2020; Moore et al., 2020b). Generally speaking, the effects of drugs after inhalation persist for a shorter time than when injected (i.p. or s.c.), and this is certainly true for heroin and THC as we’ve recently shown (Gutierrez et al., 2020a; Nguyen et al., 2016b). This difference may impact the combined effects and therefore the inhalation route was contrasted with traditional injection routes used in rodent models.

## 2. Methods

### 2.1. Subjects

Male (N=15) and female (N=7) Sprague-Dawley rats (Harlan/Envigo, Livermore, CA) and male (N=10) and female (N=22) Wistar (Charles River) rats were housed in humidity- and temperature-controlled (23±2 °C) vivaria on 12:12 hour (reversed) light:dark cycles. Rats had *ad libitum* access to food and water in their home cages and all experiments were performed in the rats’ scotophase. All procedures were conducted under protocols approved by the Institutional Care and Use Committees of The Scripps Research Institute or the University of California, San Diego.

### 2.2. Drugs

Heroin (diamorphine HCl) and Δ^9^-tetrahydrocannabinol (THC) were administered by vapor inhalation with doses described by the concentration in the propylene glycol (PG) vehicle (e.g., 50, 100 mg/mL) and duration of inhalation (15, 30 minutes) as in prior studies (Javadi-Paydar et al., 2018; Nguyen et al., 2016b). Heroin was also administered subcutaneously. THC was also administered intraperitoneally in a dose of 10 mg/kg, which produces robust temperature responses (Nguyen et al., 2016b; Taffe et al., 2015). Naloxone was administered i.p.. For injection, heroin and naloxone were dissolved in physiological saline and THC was suspended in a vehicle comprised of ethanol:cremulphor:saline in a 1:1:18 ratio. The THC and heroin were provided by the U.S. National Institute on Drug Abuse Drug Supply Program.

### 2.3. Nociception

Effects on nociception were assessed using a warm-water tail immersion assay, as described in (Javadi-Paydar et al., 2018; Nguyen et al., 2018a; Nguyen et al., 2018b). For this study, the animal’s tail was inserted ~3-4 cm into a warm (52°C) water bath and the latency for tail removal recorded with a stopwatch. A 15 second cutoff was used to avoid any potential for tissue damage.

### 2.4. Inhalation Apparatus

Vapor was delivered into sealed vapor exposure chambers (152 mm W X 178 mm H X 330 mm L; La Jolla Alcohol Research, Inc, La Jolla, CA, USA) through the use of e-vape controllers (Model SSV-3 or SVS-200; 58 watts, 0.24-0.26 ohms, 3.95-4.3 volts, ~214 °F; La Jolla Alcohol Research, Inc, La Jolla, CA, USA) to trigger Smok Baby Beast Brother TFV8 sub-ohm tanks. Tanks were equipped with V8 X-Baby M2 0.25 ohm coils. MedPC IV software was used to schedule and trigger vapor delivery (Med Associates, St. Albans, VT USA). The chamber air was vacuum-controlled by a chamber exhaust valve (i.e., a “pull” system) to flow room ambient air through an intake valve at ~1 L per minute. This also functioned to ensure that vapor entered the chamber on each device triggering event. The vapor stream was integrated with the ambient air stream once triggered. Airflow was initiated 30 seconds prior to, and discontinued 10 seconds after, each puff.

### 2.5. Radiotelemetry

Female (N=7) and male (N=15) Sprague-Dawley rats implanted with sterile radiotelemetry transmitters (Data Sciences International, St Paul, MN) in the abdominal cavity as previously described (Taffe et al., 2015; Wright et al., 2012) were used in this investigation. For studies, animals were evaluated in clean standard plastic homecages (~1cm layer of sani-chip bedding) in a dark testing room, separate from the vivarium, during the (vivarium) dark cycle. Radiotelemetry transmissions were collected via telemetry receiver plates (Data Sciences International, St Paul, MN; RPC-1 or RMC-1) placed under the cages as described in prior investigations (Aarde et al., 2013; Miller et al., 2013). Test sessions for inhalation and injection studies started with a 15-minute interval to ensure data collection, then a 15-minute interval for baseline temperature and activity values followed by the initiation of vapor sessions or drug injection.

### 2.6. Experiments

#### Experiment 1 (Inhalation Dose Ranging in Male Sprague-Dawley Rats)

A preliminary dose-ranging experiment was conducted in a group (N=8) of male Sprague Dawley rats used previously in investigations of the effects of inhalation of cannabidiol, nicotine and THC as reported (Javadi-Paydar et al., 2019a; Javadi-Paydar et al., 2019b). The first goal was to evaluate whether heroin by vapor inhalation would alter body temperature and spontaneous locomotion, as it does when injected. These were our first attempts, conducted prior to the subsequent investigations reported here and in prior publications (Gutierrez et al., 2020a; Gutierrez et al., 2020c), critical to establish efficacy of drug delivery by this method. The second goal was to identify sub-maximal exposure conditions such that interactions with another drug such as THC might be detected. All animals were habituated to the procedure with one session of 30 minutes vapor exposure to PG followed by a 60-minute recording interval. To determine the effect of heroin vapor on body temperature and locomotor activity, an initial 30 min inhalation exposure to PG and Heroin (50, 100 mg/mL in the PG) was designed. Animals were randomized to treatment conditions, and exposed, in pairs. Subsequently, the animals were evaluated after a 15-minute interval of exposure to PG, Heroin (50 mg/mL), THC (50 mg/mL) or Heroin (50 mg/mL) + THC (50 mg/mL). The treatment conditions were again counter-balanced across exposure pairs and THC was experienced no more frequently than once per week for a given rat.

#### Experiment 2 (Inhalation in Male and Female Sprague-Dawley Rats)

Groups (N=7) of male and female Sprague-Dawley rats used in studies previously reported (Gutierrez et al., 2020a) were recorded for a baseline interval, exposed to inhalation of vapor from PG, Heroin (50 mg/mL in the PG), THC (50 mg/mL) or the combination (50 mg/mL for each drug) for 30 minutes in groups of 2-3 per exposure. These studies were conducted subsequent to a series of similar challenges with vapor inhalation of nicotine, THC, and with injection of THC in additional studies not previously reported. The tail-withdrawal assay (Experiment 4, below) was conducted immediately after the vapor session and then rats were returned to their individual recording chambers for telemetry assessment. The four inhalation conditions were evaluated in a counterbalanced order across pairs/triplets, no more frequently than every 3-4 days.

#### Experiment 3 (Injection in Male and Female Sprague-Dawley Rats)

The groups (N=7) of male and female Sprague-Dawley rats used in Experiment (2 and 4) were recorded for a baseline interval, and then injected with THC (10 mg/kg, i.p.) and saline (s.c.), the 1:1:18 vehicle (i.p.) and Heroin (0.5 mg/kg, s.c.), THC (10 mg/kg, i.p.) and Heroin (0.5 mg/kg, s.c.), or the 1:1:18 vehicle and saline. For this study, treatments were twice per week (3-4 day intervals), counterbalanced across individuals, albeit with a minimum 7 day interval between any THC injections.

#### Experiment 4 (Nociception in Male and Female Sprague-Dawley Rats)

A nociception assay was conducted immediately after the vapor sessions during Experiment 2, and then rats were returned to their individual recording chambers for the telemetry assessment. The latency to withdraw the tail from insertion (3-4 cm) into a 52°C water bath was recorded by stopwatch, using techniques previously reported (Gutierrez et al., 2020c; Javadi-Paydar et al., 2018; Nguyen et al., 2019). A 15 sec cutoff was used as the maximal latency for this assay to avoid any potential for tissue damage.

#### Experiment 5 (Naloxone Antagonism in Male Sprague-Dawley Rats)

An opioid receptor antagonist study was included because we had not included this evidence in a prior study reporting heroin vapor effects on similar endpoints (Gutierrez et al., 2020a) and we’ve reported an unusual lack of effect of cannabinoid receptor 1 (CB_1_) antagonist/inverse agonist pre-treatment on THC vapor-induced hypothermia (Nguyen et al., 2020). This latter was observed despite efficacy against the anti-nociceptive effects of inhaled THC and the thermoregulatory effects in injected THC thus it was of interest to determine if a nonselective opioid receptor antagonist was effective against effects of inhaled heroin on temperature, activity and anti-nociceptive responses. The group (N=7) of male Sprague-Dawley rats used in Experiments 2-4 were recorded for a baseline interval, and then injected with either saline or naloxone (1.0 mg/kg, i.p.) 15 minutes prior to the start of a 30 minute inhalation session of either PG or Heroin (100 mg/mL) vapor. Post-inhalation, a tail-withdrawal assay was conducted (as in Experiment 4) and then animals were returned to their recording chambers for telemetric assessment. Inhalation was conducted in two pairs and one triplet, with the groupings changed on each test day. Pre-treatments were varied within the inhalation groupings to provide further randomization of conditions. Sessions were conducted two times per week (3-4 day interval) in a counterbalanced treatment order. The female group examined in parallel with these animals in prior studies was not included because the maximum number of treatment days under the approved protocol had been used in prior experiments.

#### Experiment 6 (Anti-nociception in Male and Female Wistar rats)

Groups of female (N=12) and male (N=10) Wistar rats were used in these studies. Rats had been exposed to twice daily sessions (30 minutes) of vapor from the propylene glycol (PG) vehicle or Heroin (100 mg/mL in the PG) from PND 36 to PND 45 (Gutierrez et al., 2020b). These anti-nociception experiments were conducted between PND 253 and PND 275. Four treatment conditions were evaluated in counter balanced order. Each session started with a tail-withdrawal assessment before any injections (pre-treatment). This was followed immediately with an injection of THC (10 mg/kg, i.p.) or the cannabinoid Vehicle (1:1:18 ethanol:cremulphor:saline). Another tail-withdrawal assessment was conducted 30 minutes post-injection, followed by a second injection of either Heroin (0.5 mg/kg, s.c.) or saline (the vehicle for heroin), and then a final tail-withdrawal 30 minutes later. We’ve show previously that anti-nociceptive effects of THC when injected at 5 or 10 mg/kg, i.p., persist essentially unchanged from 30 to 90 minutes after injection (Nguyen et al., 2019), justifying this sequential assessment approach. The design resulted in four replications of the pre-treatment baselines, two replications of the THC or Vehicle condition and then four final conditions of Vehicle-Saline, Vehicle-Heroin, THC-Saline and THC-Heroin.

#### Experiment 7 (Anti-nociception in Experimentally Naive Female Wistar rats)

A group (N=10) of female Wistar rats were used in these studies. Rats entered the laboratory at 11 weeks of age but due to laboratory slowdowns associated with the 2020 COVID-19 crisis remained experimentally naïve for 29 weeks. Therefore, this group was an experimentally naïve group that was well age matched with the female rats used in Experiment 5. The goal of this study was to better determine the threshold for a heroin dose which would interact with a minimal, but effective, dose of THC. Animals were first assessed for tail-withdrawal latency at baseline (no injection) or 30 minutes after heroin (0.0, 0.1, 0.32, 0.56, 1.0 mg/kg, s.c.) to better determine the lower end of the dose-effect function. For this experiment, two conditions were assessed each day with 2 h between each injection and one week between assessment days. Three within-day orders were assessed in a counterbalanced order by cage dyad, including Baseline-saline, 0.1-0.56 and 0.32-1.0. In the next experiment, rats were assessed for tail withdrawal three times in a day including a pre-treatment baseline, 30 minutes after injection of either the 1:1:18 cannabinoid vehicle or 5.0 mg/kg THC, i.p. and then 30 minutes after injection of heroin (0.0, 0.1, 0.32, 0.56 mg/kg, s.c.). Thus, the design produced 8 total treatment conditions defined by the four heroin doses and the injection of THC or its vehicle before the heroin.

### 2.7. Data Analysis

The telemeterized body temperature and activity rate (counts per minute) were collected on a 5-minute schedule in telemetry studies but are expressed as 15-minute (initial dose range study) or 30-minute averages for primary analysis. The time courses for data collection are expressed relative to the initiation of vapor inhalation and times in the figures refer to the end of the interval (e.g. “90 minutes” reflects the data collected on the six sequential 5-minute intervals from 65 to 90 minutes, inclusively). Due to transfer time after vapor sessions, the “60 minutes” interval reflects the average of collections from ~40-60 minutes). Any missing temperature values were interpolated from the values before and after the lost time point. Activity rate values were not interpolated because 5-minute to 5-minute values can change dramatically, thus there is no justification for interpolating. Data (activity rate, body temperature, tail-withdrawal latency) were generally analyzed with two-way Analysis of Variance (ANOVA) including repeated measures factors for the Drug treatment condition and the Time after vapor initiation or injection. A third factor was used for Pre-treatments, adolescent exposure group or Sex in some experiments as described in the results. A mixed-effects model was used instead of ANOVA wherever there were missing datapoints. Any significant main effects were followed with post-hoc analysis using Tukey (multi-level factors) or Sidak (two-level factors) procedures. The criterion for inferring a significant difference was set to p<0.05. All analysis used Prism 8 or 9 for Windows (v. 8.4.2 or 9.0; GraphPad Software, Inc, San Diego CA).

## 3. Results

### 3.1 Experiment 1 (Inhalation Dose Ranging in Male Sprague-Dawley Rats)

The first 30-minute inhalation duration study was originally designed to include a Heroin (100 mg/mL) condition however the first two animals randomized to this condition were substantially sedated. This condition was therefore discontinued for this experiment and these two animals were omitted from the rest of this study.

Significant effects (**Figure 1 A, C**) of Time post-initiation and the interaction of Time with Vapor condition were confirmed for both temperature [Time: F (8, 40) = 3.48 P<0.005; Interaction: F (8, 40) = 4.17; P<0.005] and activity rate [Time: F (8, 40) = 10.34; P<0.0001; Interaction: F (8, 40) = 4.48; P<0.001]. The post-hoc test confirmed that both temperature and activity were significantly lower after heroin (50 mg/mL for 30 minutes) inhalation compared with the pre-inhalation baseline and with the PG inhalation condition in the interval 45-60 minutes after the start of inhalation.

**Figure 1:**
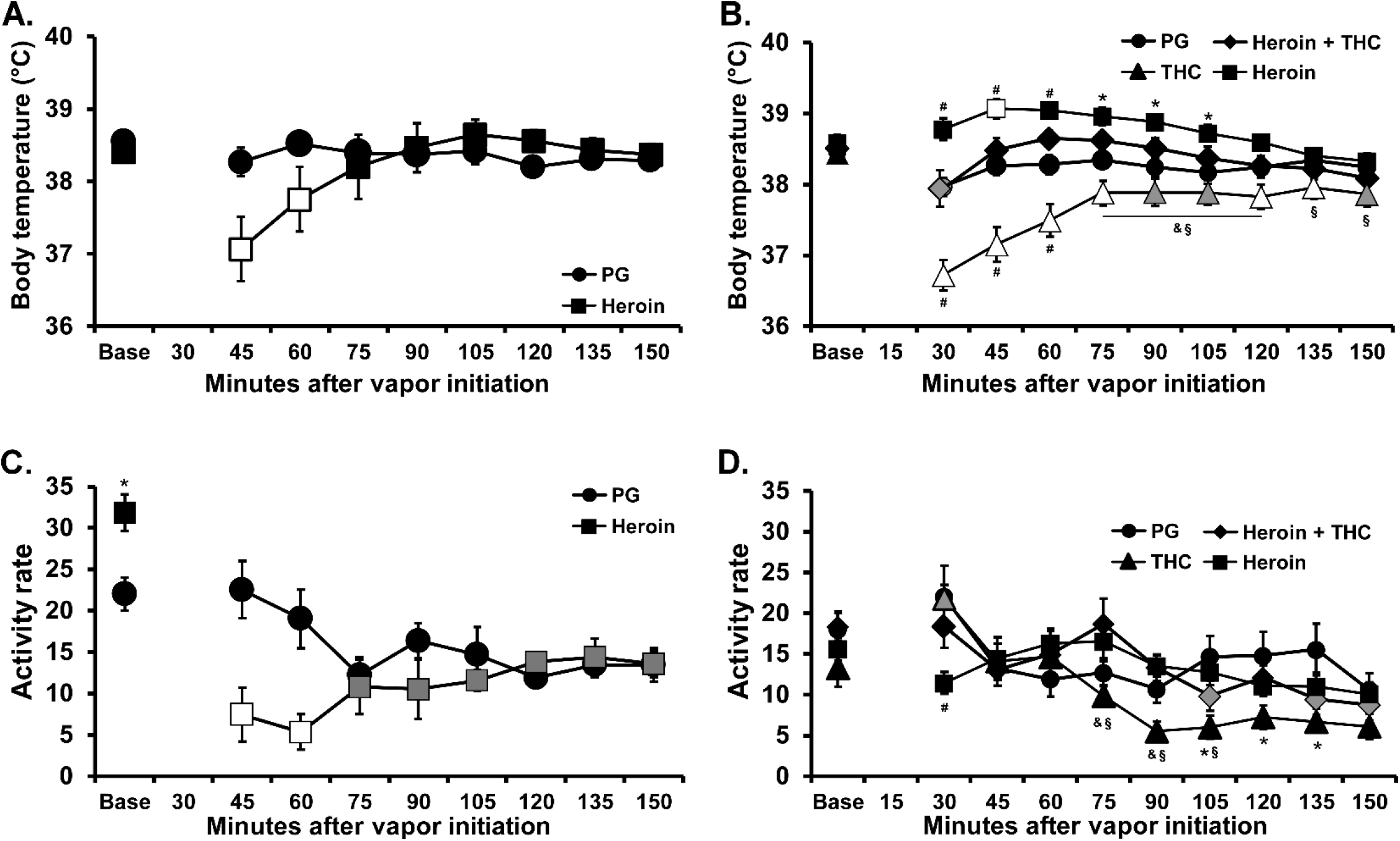
Mean A) body temperature and C) activity rate for male Sprague-Dawley rats (N=6; ±SEM) after vapor inhalation of the propylene glycol vehicle (PG) or Heroin (50 mg/mL in the PG) for 30 minutes. Mean B) body temperature and D) activity rate for male Sprague-Dawley rats (N=8; ±SEM) after vapor inhalation of the propylene glycol vehicle (PG), Heroin (50 mg/mL in the PG), THC (50 mg/mL) or the combination (Heroin + THC) for 15 minutes. Open symbols indicate a significant difference from both the vehicle at a given time-point and the within-treatment baseline, while shaded symbols indicate a significant difference from the baseline, within treatment condition, only. A significant difference from the PG inhalation condition is indicated by *, a difference from the Heroin condition with §, a difference from the Heroin + THC conditions with &, and from all other conditions with #.

In the 15 minute exposure study (**Figure 1 B, D**), the ANOVAs confirmed that there were significant effects of Time post-initiation [F (9, 63) = 11.04; P<0.0001], Vapor condition [F (3, 21) = 17.75; P<0.0001] and the interaction of Time with Vapor condition [F (27, 189) = 7.09; P<0.0001] on temperature and significant effects of Time [F (9, 63) = 11.60; P<0.0001] and the interaction of Time with Vapor condition [F (27, 189) = 2.60; P<0.0001] on activity rate. The post hoc test further confirmed that body temperature was higher after heroin inhalation (30-105 minutes after the start of exposure), and lower in the THC condition (30-105, 120-135 minutes), compared with the PG condition; temperature likewise differed between THC and Heroin conditions (30-150 minutes). Temperature was also higher in the heroin condition (30-60 minutes) and lower in the THC condition (30-120 minutes) in comparison with temperature in the THC+Heroin combined condition. The post hoc exploration also confirmed that activity was lower in the heroin (30 minutes) and THC conditions (105-135 minutes) compared with the PG condition. Significantly lower activity was also confirmed relative to the THC+Heroin condition for heroin (30 minutes) and THC (75-90 minutes) and activity differed between THC and Heroin conditions (30, 75-105 minutes).

### 3.2 Experiment 2 (Inhalation in Male and Female Sprague-Dawley Rats)

The body temperature of the female rats (**Figure 2 A**) was significantly elevated by the inhalation of Heroin 50 mg/mL and decreased by the inhalation of THC (50 mg/mL). The ANOVA confirmed a significant effect of Time after vapor initiation [F (8, 48) = 25.65; P<0.0001], of Vapor Condition [F (3, 18) = 43.71; P<0.0001] and of the interaction of Time with Vapor Condition [F (24, 144) = 16.13; P<0.0001] on body temperature. The post-hoc test confirmed that temperature was significantly higher after inhalation of Heroin 50 mg/mL (90-120 minutes after the start of inhalation), and lower after inhalation of THC 50 mg/mL (60-270 minutes) compared with the PG condition. Temperature following the inhalation of both drugs in combination was significantly lower compared with PG (60-90, 210-270 minutes), Heroin (60-210 minutes) and higher compared with THC (60-180 minutes). Temperature was also significantly lower than the baseline observation 60-270 minutes after the inhalation of THC or the combination, but not in the other two vapor conditions.

**Figure 2:**
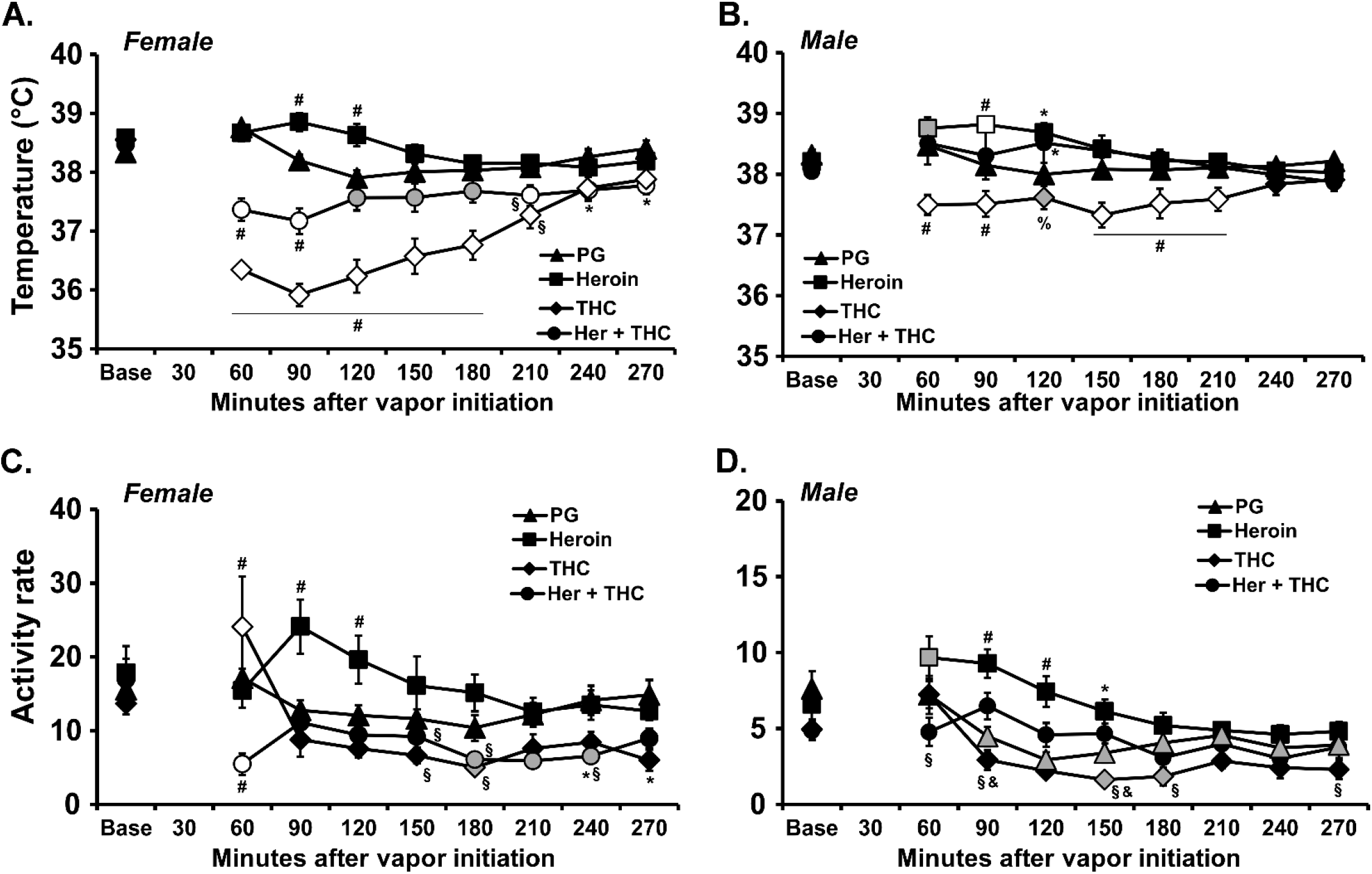
Mean (N=7 per group; ±SEM) body temperature (A, B) and activity rate (C, D) for female (A, C) and male (B, D) Sprague-Dawley rats after vapor inhalation of the propylene glycol vehicle (PG), <u>Her</u>oin (50 mg/mL in the PG), THC (50 mg/mL) or the combination of Heroin and THC (at 50 mg/mL each) for 30 minutes. N.b. there is a two fold difference in the scale of the activity in C and D, to accommodate the sex difference in activity rate. Open symbols indicate a significant difference from both the vehicle at a given time-point and the within-treatment baseline, while shaded symbols indicate a significant difference from the baseline, within treatment condition, only. A significant difference from the PG inhalation condition is indicated by *, a difference from the Heroin condition with §, a difference from the THC condition with %, a difference from the Heroin + THC conditions with &, and from all other conditions with #.

The activity of the female rats (**Figure 2 C**) was increased after heroin inhalation and decreased after inhalation of the THC+Heroin combination. The ANOVA confirmed a significant effect of Time after vapor initiation [F (8, 48) = 9.81; P<0.0001], of Vapor Condition [F (3, 18) = 6.97; P=0.0026] and of the interaction of Time with Vapor Condition [F (24, 144) = 3.91; P<0.0001] on activity rate. The post-hoc test confirmed that activity was decreased relative to the baseline and all other inhalation conditions 60 minutes after the start of THC+Heroin inhalation and elevated relative to the baseline and all other inhalation conditions 60 minutes after the start of THC. Activity was elevated compared with all other conditions 90-120 minutes after the start of Heroin inhalation.

The body temperature of the male rats (**Figure 2 B**) was also significantly elevated by the inhalation of Heroin 50 mg/mL and decreased by the inhalation of THC (50 mg/mL). The ANOVA confirmed a significant effect of Time after vapor initiation [F (8, 48) = 2.64; P<0.05], of Vapor Condition [F (3, 18) = 7.84; P=0.0015] and of the interaction of Time with Vapor Condition [F (24, 144) = 5.26; P<0.0001] on body temperature. Temperature was significantly lowered after the inhalation of THC (60-210 minutes after the start of inhalation) and was significantly increased 60-90 minutes after the start of Heroin inhalation, relative to the Baseline observation, but was unchanged in the other two conditions. Similarly, the post-hoc test confirmed that temperature was significantly higher after inhalation of Heroin 50 mg/mL (90-120 minutes after the start of inhalation), and lower after inhalation of THC 50 mg/mL (60-90, 150-210 minutes) or the combination (120 minutes), compared with the PG condition. Temperature following the inhalation of both drugs in combination was significantly lower compared with Heroin (90 minutes) and higher compared with THC (60-210 minutes).

The activity of the male rats was (**Figure 2D**) was elevated after heroin inhalation, as confirmed with a significant effect of Time after vapor initiation [F (8, 48) = 13.41; P<0.0001], of Vapor Condition [F (3, 18) = 16.81; P<0.0001] and of the interaction of Time with Vapor Condition [F (24, 144) = 2.69; P=0.0001] on activity rate in the ANOVA. The post-hoc test confirmed that activity was elevated 90-120 minutes after the start of Heroin inhalation compared with all other conditions, as well as at individual timepoints relative to the combination (60 minutes), THC (150, 180, 270 minutes), and PG (150 minutes). Activity differed between TH and the combined inhalation (90, 150 minutes), was higher than the baseline 60 min after initiation of heroin and lower than the baseline in the PG (90-270 minutes after initiation) and THC (150-180 minutes) inhalation conditions.

### 3.3 Experiment 3: (Injection in Male and Female Sprague-Dawley Rats)

The injection of THC (10 mg/kg, i.p.) and heroin (0.5 mg/kg, s.c.) produced opposing effects on activity in male and female groups, and opposing effects on temperature in the male rats (**Figure 3**). The body temperature of the female rats was significantly decreased by THC (**Figure 3 A**). The ANOVA confirmed a significant effect of Time after injection [F (9, 45) = 13.44; P<0.0001], of Drug Treatment Condition [F (3, 15) = 16.13; P<0.0001] and of the interaction of Time with Drug Condition [F (27, 135) = 7.14; P<0.0001] on body temperature in the female rats. The post-hoc test confirmed that temperature was changed relative to the baseline, and also relative to the ehicle+Saline and Heroin injection conditions, after the injection of THC (30-270 minutes post-injection) or the combination (30-270 minutes).

**Figure 3:**
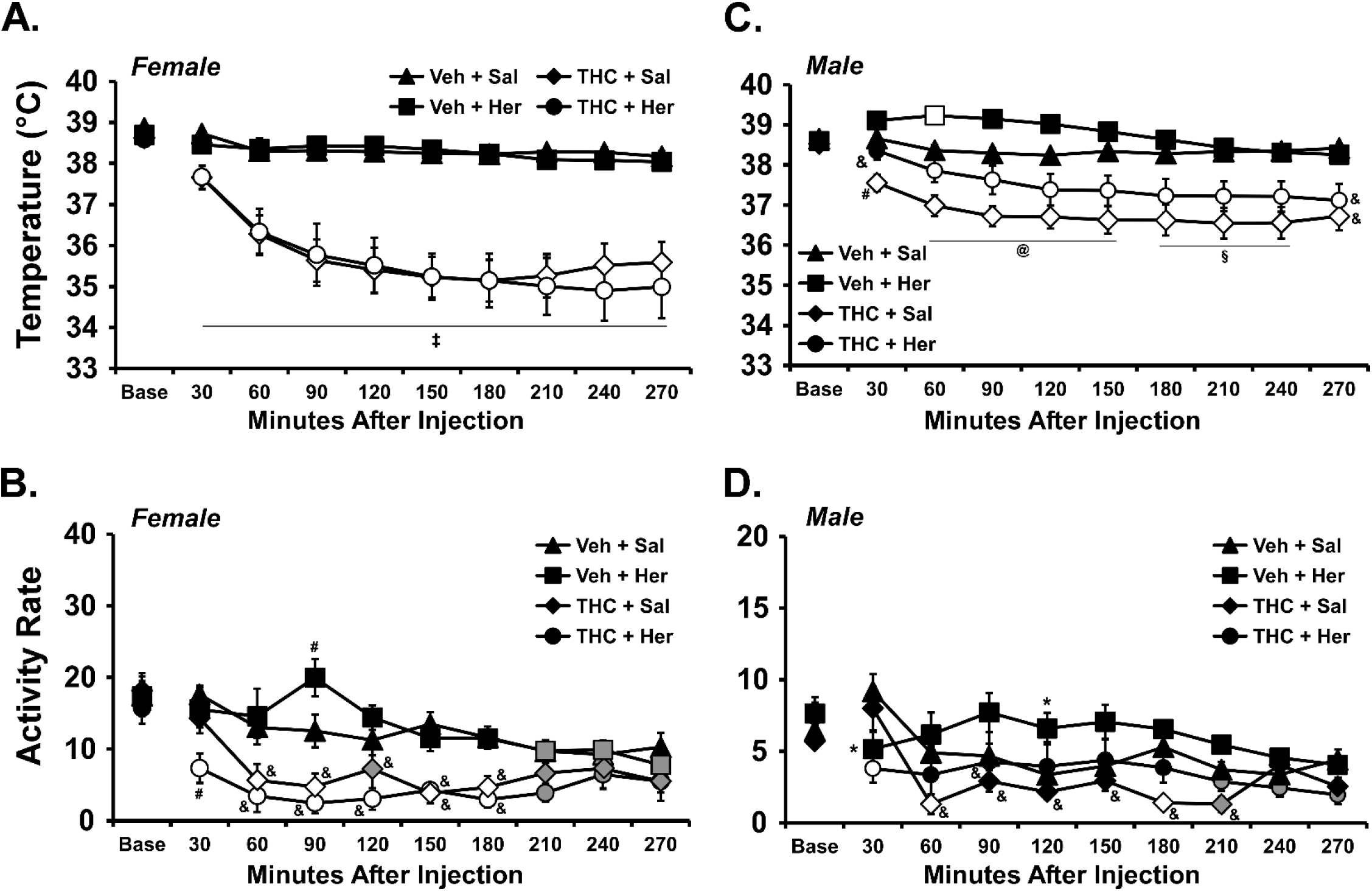
Mean (±SEM) body temperature (A, C) and activity rate (B, D) for female (N=7; A, B) and male (N=6; C,D) Sprague-Dawley rats after injection of THC (10.0 mg/kg, i.p.), Heroin (0.5 mg/kg, s.c.), the 1:1:18 <u>Veh</u>icle or <u>Sal</u>ine in the indicated combinations. N.b. there is a two fold difference in the scale of the activity in B and D, to accommodate the sex difference in activity rate. Open symbols indicate a significant difference from both the vehicle at a given time-point and the within-treatment baseline, while shaded symbols indicate a significant difference from the baseline, within treatment condition, only. A significant difference of all treatment conditions from each other at a given time post-injection is indicated with @, a difference between all active drug conditions with § and a difference of each of the THC conditions from both of the Vehicle conditions with ‡. A significant difference from the PG inhalation condition (but not the baseline) is indicated by *, a difference from the Heroin condition with &, and from all other conditions with #.

The activity of the female rats was increased by heroin injection and decreased by the THC injection (**Figure 3 B**) as confirmed by a significant effect of Time after injection [F (9, 54) = 9.47; P<0.0001], of Drug Treatment Condition [F (3, 18) = 17.61; P<0.0001] and of the interaction of Time with Drug Condition [F (27, 162) = 2.43; P<0.0005] on activity in the ANOVA. The post-hoc test confirmed that activity rates were reduced compared with the baseline and the Vehicle+Saline injection condition after THC+Saline (60-90, 150-180 minutes) or THC+Heroin (60-180 minutes), reduced compared with all other conditions 30 minutes after injection of THC+Heroin and increased compared with all other conditions 90 minutes after injection of Vehicle+Heroin. Significantly lower activity compared with Heroin alone was also observed in the THC+Heroin condition (30-180 minutes after the start of inhalation) as well as the THC condition (60-180 minutes).

The body temperature of the male rats was significantly elevated by heroin and decreased by THC (**Figure 3 C**). The ANOVA confirmed a significant effect of Time after injection [F (9, 54) = 26.79; P<0.0001], of Drug Treatment Condition [F (3, 18) = 15.12; P<0.0001] and of the interaction of Time with Drug Condition [F (27, 162) = 6.26; P<0.0001] on body temperature. The post-hoc test confirmed that temperature was changed relative to the baseline, and also relative to the vehicle/saline injection condition, after the injection of THC (30-270 minutes post-injection), Heroin (60 minutes) or the THC + Heroin combination (60-270 minutes). Body temperature was also significantly lower compared with the Heroin condition after the injection of either THC (30-270) or the THC+Heroin combination (30-270 minutes).

The activity of the male rats (**Figure 3 D**) was increased by heroin injection and decreased by the THC injection as confirmed by a significant effect of Time after injection [F (9, 54) = 6.27; P<0.0001], of Drug Treatment Condition [F (3, 18) = 9.20; P=0.001] and of the interaction of Time with Drug Condition [F (27, 162) = 2.34; P<0.001] on activity in the ANOVA. The post-hoc test confirmed that activity rates were reduced compared with the baseline and the Vehicle+Saline injection condition after THC+Saline (60, 180 minutes after injection) or THC+Heroin (30 minutes), and increased relative to the Vehicle+Saline injection condition after Vehicle+Heroin (120 minutes after injection).

### 3.4 Experiment 4 (Nociception in Male and Female Sprague-Dawley Rats)

The ANOVA confirmed that Sex [F (1, 12) = 10.10; P<0.01] and Vapor Condition [F (4, 48) = 27.94; P<0.0001] significantly altered the tail withdrawal latency (**Figure 4**). The post-hoc test failed to confirm any specific sex differences within any of the Vapor conditions. Within the female or male groups, a significant difference from the PG condition was observed after the inhalation of Heroin or the combination of Heroin and THC. Tail withdrawal was likewise significantly slower after the inhalation of the combination relative to the THC inhalation in the female rats and relative to the THC or Heroin conditions in the male rats.

**Figure 4:**
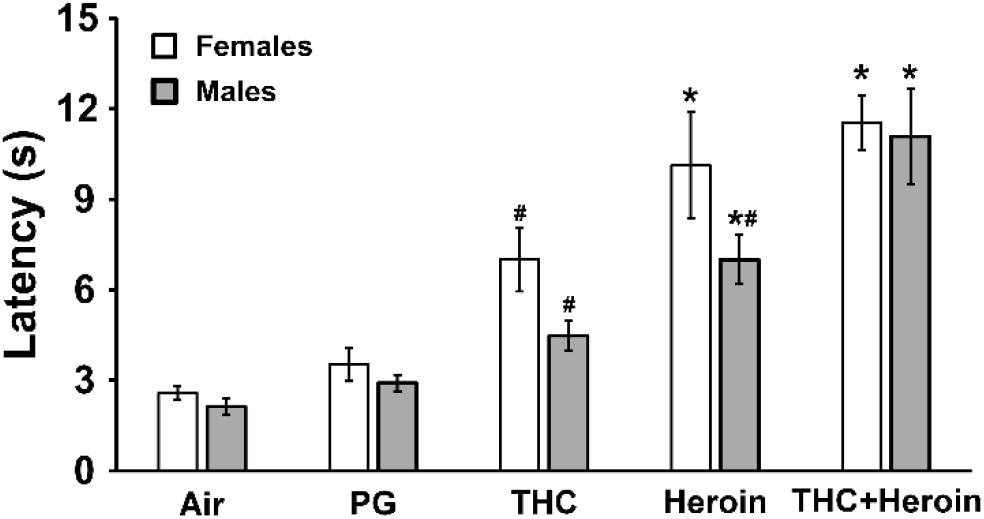
Mean (±SEM) tail withdrawal latency for male and female (N=7 per group) rats after vapor exposure. A significant difference from PG condition is indicated with * and a difference from the THC+Heroin condition with #. PG, propylene glycol. Heroin (50 mg/mL); THC (50 mg/mL).

### 3.5 Experiment 5 (Naloxone Antagonism in Male Sprague-Dawley Rats)

#### 3.5.1 Temperature and Activity

Pre-treatment with naloxone attenuated the temperature and locomotor responses to heroin inhalation (**Figure 5**). One individual exhibited an anomalous temperature response to each heroin condition and was omitted from the final analyses, thus N=6 for this study. As in Experiment 2, after inhaling the vehicle vapor the body temperature of rats declined slightly across the first three hours, and the prior injection of naloxone had no impact. The inhalation of heroin (100 mg/mL) for 30 minutes resulted in higher body temperature in the first two hours after the start of inhalation, compared with the PG inhalation conditions (**Figure 5 A**); again, this was similar to the outcome of heroin inhalation in Experiment 2. Naloxone pre-treatment attenuated the temperature response to heroin. The three-way ANOVA confirmed a significant effect of the interaction of Vapor Condition with Time after the start of inhalation [F (9, 45) = 15.45; P<0.0001], of the interaction of Pre-treatment Condition with Time after the start of inhalation [F (9, 45) = 2.90; P<0.01] and the interaction of all three factors [F (9, 45) = 3.57; P<0.005] on body temperature. The post-hoc test confirmed first that heroin inhalation resulted in increased body temperature relative to PG inhalation 90-180 minutes after initiation. In addition, while naloxone had no impact on temperature when administered prior to PG inhalation, it attenuated the increase in body temperature observed 90-150 minutes after the start of Heroin inhalation. Activity was also impacted by heroin inhalation and the three-way ANOVA confirmed a significant effect of Time after the start of inhalation [F (9, 45) = 8.60; P<0.0001], of Vapor Condition [F (1, 5) = 7.17; P<0.05], of the interaction of Vapor Condition with Time after the start of inhalation [F (9, 45) = 5.09; P<0.0001] on activity. The post-hoc test confirmed that naloxone pre-treatment caused significant attenuation of heroin-stimulated activity 90 minutes after the start of inhalation.

**Figure 5:**
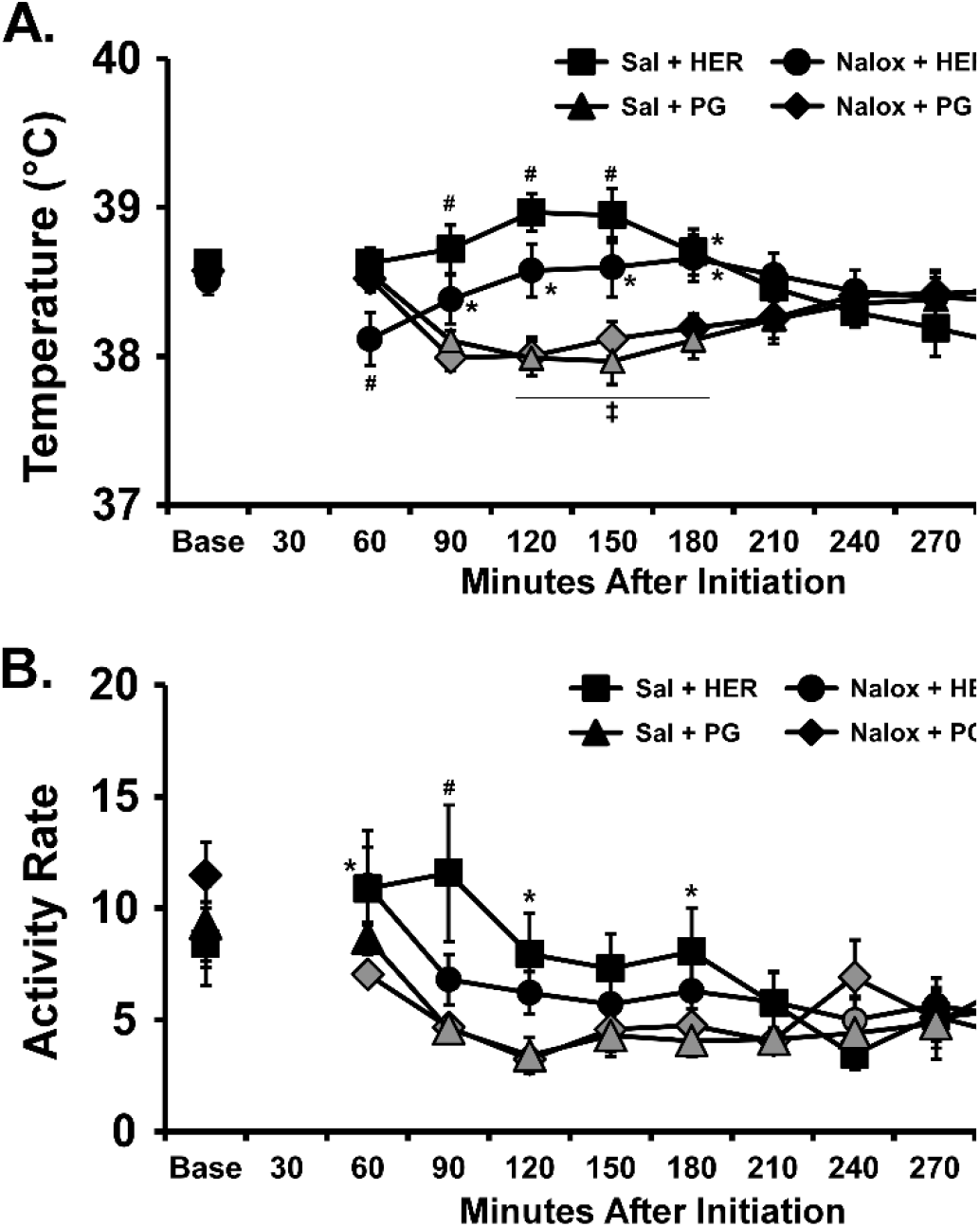
Mean (N=7; ±SEM) A) temperature and B) Activity for male Sprague-Dawley rats after injection with either <u>Sal</u>ine or <u>Nalox</u>one (1.0 mg/kg, i.p.) and propylene glycol (PG) or Heroin (HER; 100 mg/mL) vapor exposure for 30 minutes. Shaded symbols indicate a significant difference from the <u>Base</u>line, within treatment condition. A significant difference of each of the heroin vapor conditions from each of the PG conditions is indicated with ‡. A significant difference from the PG inhalation condition (within pre-treatment) is indicated by *, and a difference compared with all other conditions with #.

#### 3.5.2 Anti-nociception

The tail withdrawal latency was increased by Heroin vapor inhalation (**Figure 6**) and this effect was attenuated by the pre-inhalation injection of naloxone. The ANOVA confirmed significant effects of Vapor Condition [F (1, 6) = 20.18; P<0.005] and of the interaction of Pre-treatment and Vapor conditions [F (1, 6) = 6.69; P<0.05], but not of Pre-treatment alone [F (1, 6) = 5.94; P=0.051]), on tail withdrawal latency. The post-hoc test confirmed that this was attributable to tail-withdrawal latency being longer in the Saline-Heroin vapor condition compared with the other three treatment conditions.

**Figure 6:**
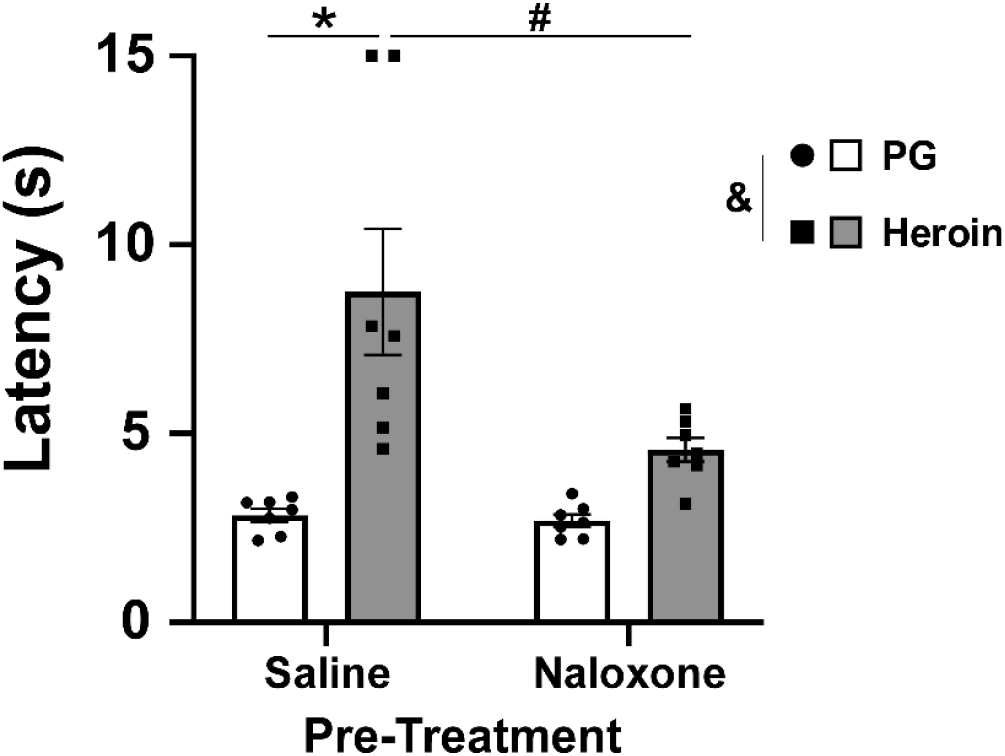
Mean (±SEM) tail withdrawal latency for male (N=7) Sprague-Dawley rats after 30 minutes of heroin vapor exposure, following injection of saline or naloxone (1.0 mg/kg, i.p.). A significant difference across inhalation conditions is indicated with &, a significant difference from PG inhalation, within pre-treatment condition, is indicated with * and a difference from the Saline pre-treatment within inhalation condition with #. PG, propylene glycol. Heroin (100 mg/mL).

### 3.6 Experiment 6 (Nociception in Male and Female Wistar Rats)

For analysis, each of the three tail-withdrawal assays on each of four treatment sequence days was considered as a unique Observation Condition (**Figure 7**). This resulted in four pre-treatment observations, two after the 1:1:18 vehicle injection, two after THC (10 mg/kg, i.p.) injection and one and the end of each of the Vehicle+Saline, Vehicle+Heroin, THC+Saline and THC+Heroin sequences. The initial three-way ANOVA confirmed significant effects of Sex [F (1, 216) = 13.75; P<0.0005], of the Observation Condition [F (11, 216) = 39.87; P<0.0001] and an interaction of Sex with Adolescent Treatment [F (1, 216) = 4.29; P<0.05] on tail-withdrawal latency in the Wistar rats. The first follow-up two-way ANOVA collapsed across adolescent treatment (**Figure 7 B**) confirmed a significant effect of Observation Condition [F (11, 220) = 49.85; P<0.0001] and of Sex [F (1, 20) = 4.66; P<0.05]. The post-hoc test of the marginal means for treatment condition confirmed that tail-withdrawal latency was higher after each of the THC injections compared with all of the pre-treatment latencies and with the two latencies after the 1:1:18 Vehicle injections. It was further confirmed that latencies were higher after the Heroin injection in the Vehicle condition and after the saline injection in the THC condition compared with the respective pre-injection baselines and with the latency in the Vehicle+Saline condition. Finally, tail-withdrawal latencies after the heroin injection in the THC condition were significantly higher than in any other condition. Within sex, these patterns were similarly confirmed for the female group. Within the male subgroup, withdrawal latencies were higher after the THC+Heroin condition compared with all other treatment conditions and in the THC+Saline condition compared with the respective pre-treatment baseline and the Vehicle+Saline condition. The only sex difference confirmed within a specific treatment condition was after injection of both THC and Heroin. Collapsed across sex, there was no significant effect of adolescent exposure group, an effect of Observation condition [F (11, 220) = 48.81; P<0.0001] and the posthoc test recapitulated the overall outcome (**Figure 7 C**).

**Figure 7:**
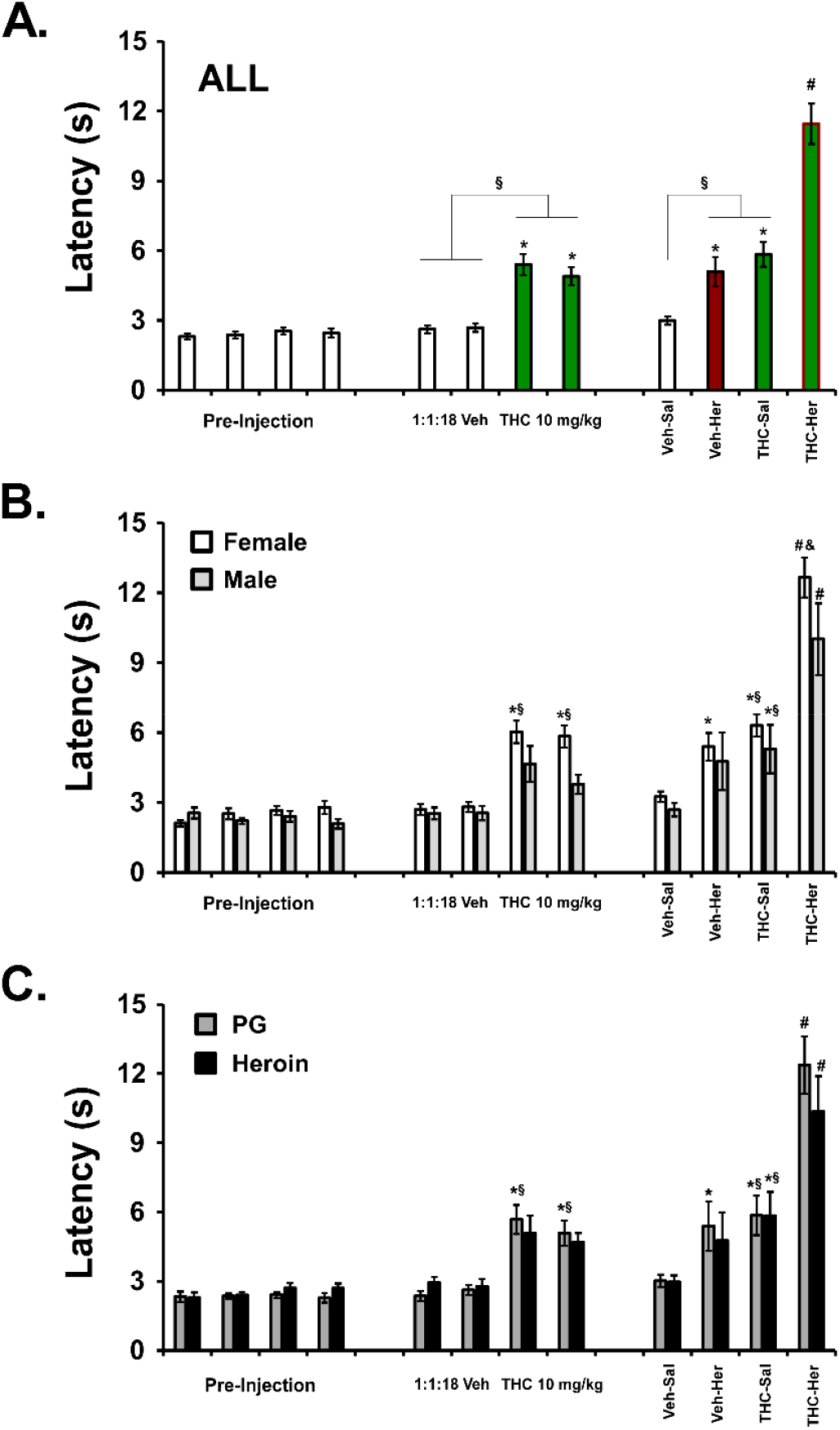
A) Mean (±SEM) tail withdrawal latency for all rats (N=22) at three timepoints within each of four treatment conditions/sessions. The first injection of the session was either THC (10 mg/kg, i.p.) or the cannabinoid vehicle and the second injection was either saline or Heroin (0.5 mg/kg, s.c.) for four total treatment conditions. B) Withdrawal latency for female (N=12) and male (N=10) subgroups. C) Withdrawal latency for subgroups of rats treated with repeated PG (N=12) or Heroin (N=10) vapor as adolescents. Within groups, a significant difference from the respective pre-injection baseline is indicated with *, a difference from the respective Vehicle or Saline condition with §, a difference from all other treatment conditions with #. A difference between groups is indicated with &.

### 3.7 Experiment 7 (Nociception in Experimentally Naive Female Wistar Rats)

The injection of heroin (0.1-1.0 mg/kg, s.c.) by itself produced dose-dependent increases in tail-withdrawal in the female Wistar rats (N=10), as shown in **Figure 8 A.**The ANOVA confirmed a significant effect of dose [F (2.031, 18.28) = 31.89; P<0.0001], and the post-hoc test confirmed that withdrawal latency was longer after 0.56 mg/kg, compared with the vehicle injection, and longer after 1.0 mg/kg compared with all other conditions. Using the threshold doses of heroin, it was then shown that interactive effects with THC (5 mg/kg, i.p.) were only observed at the 0.56 mg/kg dose of heroin (**Figure 8 B, C**). Analysis included a factor for the within-session time of observation as well as the final heroin dose condition. The two-way ANOVA within the *vehicle pre-treatment condition* first confirmed a significant effect of the within-session assessment time [F (2, 18) = 24.42; P<0.0001], but not of Heroin dose condition or the interaction of the factors, on withdrawal latency (**Figure 8 B**). The post-hoc confirmed that this was attributable to a significant increase compared with the pre-treatment assessment after the vehicle injection for the Heroin 0.1 mg/kg treatment, and a difference of the post-Heroin 0.56 mg/kg latency compared with the pretreatment and post-vehicle assessment on that same session. In contrast, the two-way ANOVA within the *THC 5.0 mg/kg, i.p., pre-treatment condition* confirmed an effect of the within-session assessment time [F (2, 18) = 77.73; P<0.0001], of Heroin dose condition [F (3, 27) = 10.96; P<0.0001] and of the interaction of the factors [F (6, 54) = 5.14; P<0.0005] on tail withdrawal latency (**Figure 8 C**). The post-hoc test first confirmed that the effect of heroin dose (on the final assessment of the session) was attributable to a significant increase of latency in all three active dose conditions compared with saline, and an elevation after heroin 0.56 mg/kg compared with the 0.1 or 0.32 mg/kg conditions. The post-hoc test further confirmed that the THC injections increased tail-withdrawal latency compared with the pre-treatment assessment within all four sessions. Heroin only significantly increased latency beyond the effect of the THC injection in the 0.56 mg/kg condition. Most critically, a two-way ANOVA analyzing the withdrawal latency after the last observation of all sessions (i.e., after heroin injection in both THC and vehicle pre-treatment conditions) confirmed a significant effect of the Heroin dose [F (3, 27) = 11.46; P<0.0001], of the Vehicle/THC pre-treatment [F (1, 9) = 46.94; P<0.0001] and of the interaction of factors [F (3, 27) = 4.30; P<0.05] on tail-withdrawal latency. The post-hoc test further confirmed a significant difference between saline injection and all three heroin doses, and between 0.32 and 0.56 mg/kg, within the THC pre-treatment condition. There were no effects of heroin dose in the vehicle pretreatment conditions confirmed in this analysis.

**Figure 8:**
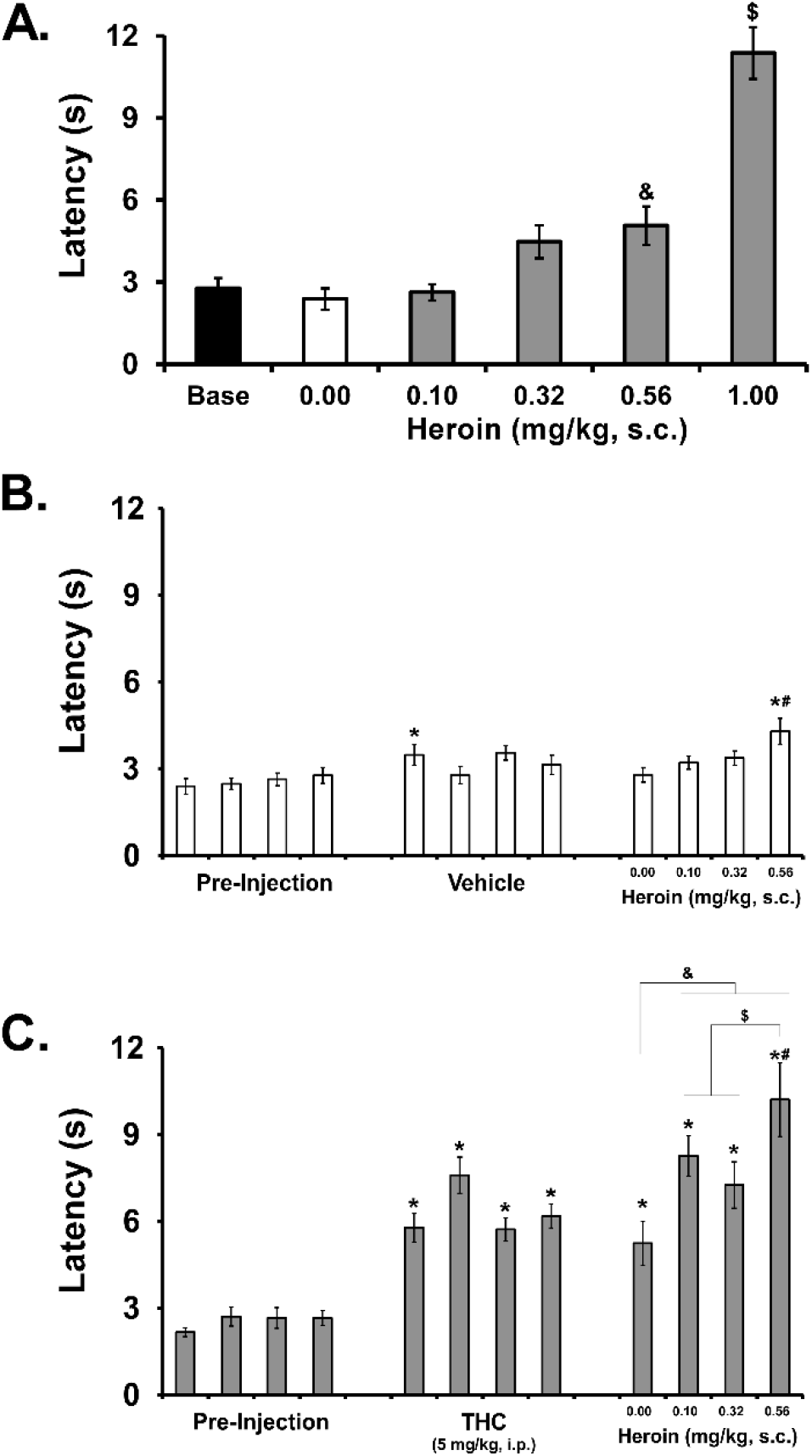
A) Mean (±SEM) tail-withdrawal latency in a group (N=10) of female Wistar rats evaluated after no treatment (<u>Base</u>line) or after injection with saline or heroin (0.1-0.56 mg/kg, s.c.). Mean (±SEM) tail-withdrawal latency in a group (N=10) of female Wistar rats evaluated before injection, then after either B) the 1:1:18 vehicle or C) THC 5 mg/kg, i.p., then after heroin. A significant difference from the saline condition is indicated by & and a difference from all other conditions with $. Within-session, a significant difference from the pre-injection value is indicated by * and a difference from the vehicle/THC injection with #. Across sessions, a significant difference from the saline (0.0) condition is indicated with & and from the 0.56 dose with $.

## 4. Discussion

This study first confirms that THC enhances the anti-nociceptive effect of heroin when the drugs were administered by vapor inhalation through the use of an Electronic Drug Delivery System (EDDS) (**Figure 4**), similar to effects observed when delivered by parenteral injection (**Figures 7, 8**). This outcome is consistent with a prior demonstration that THC enhances the effects of another opioid, oxycodone, on anti-nociception when drugs are delivered either by inhalation or injection (Nguyen et al., 2019). Thus, the routes of administration and the doses used in this study are those which are effective at demonstrating the “opioid sparing” effects of THC when it comes to one desired medical benefit of both opioids and THC, i.e., analgesia.

In contrast, the effects of each drug on thermoregulation and locomotor activity when co-administered appear to be independent. That is, when doses of heroin that increase body temperature are combined with doses of THC that decrease body temperature the net effect is an intermediate response, indistinguishable from the response to vehicle in some cases. Interpretation of the effects of co-administration on activity are complex because the effects of higher doses of heroin are biphasic with time after dosing. Still, at a time post-administration when <u>*hyper*</u>locomotive effects of heroin are observed, the combination with <u>*hypo*</u>locomotor doses of THC result in an intermediate effect on activity.

There is a biphasic dose-dependent effect of heroin on body temperature with lower exposures producing small, but reliable, increases in temperature and higher doses / exposures reducing body temperature, as with the first experiment (**Figure 1**) and in female rats in a prior report (Gutierrez et al., 2020a). Here, we selected parameters of exposure to heroin anticipated to increase body temperature in both male and female rats and a parameters of THC exposure anticipated to lower body temperature. The results show (**Figure 1 B, Figure 2 A, B**) that the co-administration produces an intermediate phenotype, consistent with independent effects. This was the case in two cohorts of male rats as well as in the female rats, illustrating robustness of the observation. There was a slight disconnect in that inhalation of vapor from Heroin 50 mg/mL for 30 minutes was the higher exposure in the first group and constituted a lower (phenotypic) exposure in the second group of males. It is likely that the slight change in methods across studies (and in particular the e-cigarette cannister type) produced the difference, further underlining the necessity for validation studies to hone exposure conditions when using this method.

We have previously shown that heroin vapor exposure (100 mg/mL, 30 minutes) produces anti-nociceptive effects in male and female Wistar rats (Gutierrez et al., 2020c) and that is herein extended to Sprague-Dawley rats. We also show here that the competitive, non-selective opioid receptor antagonist naloxone attenuates the anti-nociceptive effects of inhaled heroin (**Figure 6**). It also blunted the thermoregulatory and locomotor stimulant responses to heroin inhalation (**Figure 5**). This is a novel observation since we did not include this in our prior study (Gutierrez et al., 2020a) which was the first to show efficacy of EDDS vapor delivery of heroin. While the effect of naloxone is perhaps expected, we’ve reported an unanticipated lack of effect of CB_1_ antagonist/inverse agonist pre-treatment on THC vapor-induced hypothermia (Nguyen et al., 2020), so it was important to confirm opioid antagonist efficacy in this model. The failure to achieve a statistically reliable additive anti-nociceptive effect of heroin inhalation in the female group is likely due to a slightly more robust response to the heroin inhalation condition, compared with the males. We then went on to expand this experiment to a broader range of conditions in female Wistar rats, using drug injection to afford tighter dose control (**Figure 8**). These studies demonstrated that a 0.56 mg/kg, s.c., dose of heroin combined with a 5.0 mg/kg, i.p. dose of THC appears to approximate the threshold for observable interactions. After demonstrating that the 0.1-0.32 mg/kg, s.c., doses of heroin produce minimal effect administered alone, we then went on to show in the complex design that only 0.56 mg/kg produced a significant increase in nociception beyond that induced by 5.0 mg/kg, i.p., THC. The 5.0 and 10.0 mg/kg THC doses appear to produce approximately the same magnitude of effect in the tail withdrawal assay at 52°C as illustrated here (**Figure 7, 8**) and in prior work (Nguyen et al., 2016b; Nguyen et al., 2019), suggesting an asymptote in the dose-response curve for this drug when administered alone. It is therefore intriguing that the addition of 0.5 mg/kg heroin to 10 mg/kg THC had greater additive anti-nociceptive effect than the addition of 0.56 mg/kg heroin to 5 mg/kg THC. This hints at a more than additive interactive effect that would need to be explored with additional experimental designs to confirm. Overall, these results confirm the so-called opioid-sparing effects of THC in the context of thermal analgesia. The translational potential is that by taking lower doses of each drug the consequences of higher doses might be partially avoided. This may be important for slowing the development of tolerance to the therapeutic benefits of either drug independently and thereby slow the need for increasing doses.

In a more general sense, this study further confirms the utility of the EDDS approach for the investigation of poly-substance use. Patterns of poly drug use in humans via simultaneous vaping are already being described in the epidemiological literature (Dunbar et al., 2020), which tends to lag real world practices by months to years, thus it is a critical advance to be able to study such practices in a controlled laboratory model. As outlined in the Introduction, EDDS models for the inhalation delivery of a range of drugs in laboratory rodents are being rapidly developed and reported. This technique for laboratory rodent research is flexible in terms of drug substances and doses and is therefore capable of supporting poly-drug investigations. We have previously used this model to explore interactive effects of nicotine with THC (Javadi-Paydar et al., 2019b) and of nicotine with cannabidiol (Javadi-Paydar et al., 2019a), as well as the effects of combined inhalation of these two cannabinoids (Javadi-Paydar et al., 2018).

In conclusion, THC can reduce the dose of heroin necessary for a given analgesic effect. However, the effects of combined THC and heroin on activity and thermoregulation show, critically, that the inference that THC *universally* enhances the effects of heroin, or vice versa, is not supported. When the effects of each drug in isolation are in the same direction, due to either dose or the *in vivo* endpoint, they can appear to have interactive effects. However, when the effects are in the opposite direction, such as with body temperature and activity at specific doses, then the combination produces an intermediate phenotypic outcome. The anti-nociception data do not suggest any super-additive interactive effects of THC with heroin. While a super-additive interaction might appear beneficial for medical applications the limitation would be that any tolerance to one or the other drug that appears would also have interactive, rather than merely subtractive effects. Additional study would be required to determine whether combinations of cannabinoid and opioid drugs used chronically result in less rapid tolerance compared with equipotent therapeutic regimens of each drug taken alone. Relatedly, it remains to be determined if the ability to more closely titrate dose to effect can be obtained with the inhaled route of administration of THC or an opioid.

## Acknowledgements

The study was conducted with the support of USPHS grants (R01 DA035281, R01 DA042211), a UCSD Chancellor’s Post-doctoral Fellowship (AG) and the Tobacco Related-Disease Research Program (TRDRP; T31IP1832). The NIH/NIDA and the TRDRP had no role in study design, collection, analysis and interpretation of data, in the writing of the report, or in the decision to submit the paper for publication. The authors declare no additional financial conflicts which affected the conduct of this work.

## Literature Cited

Aarde, S.M., Huang, P.K., Creehan, K.M., Dickerson, T.J., Taffe, M.A., 2013. The novel recreational drug 3,4-methylenedioxypyrovalerone (MDPV) is a potent psychomotor stimulant: self-administration and locomotor activity in rats. Neuropharmacology 71, 130–140.

Ahmad, T., Lauzon, N.M., de Jaeger, X., Laviolette, S.R., 2013. Cannabinoid transmission in the prelimbic cortex bidirectionally controls opiate reward and aversion signaling through dissociable kappa versus mu-opiate receptor dependent mechanisms. J Neurosci 33(39), 15642–15651.

Allem, J.P., Majmundar, A., Dharmapuri, L., Cruz, T.B., Unger, J.B., 2019. E-liquid-related posts to Twitter in 2018: Thematic analysis. Addict Behav Rep 10, 100196.

Blakesley, B.C., Dinneen, L.C., Elliott, R.D., Francis, D.L., 1972. Intravenous self-administration of heroin in the rat: experimental technique and computer analysis. Br J Pharmacol 45(1), 181P–182P.

Breit, K.R., Rodriguez, C., Lei, A., Thomas, J.D., 2020. Combined Vapor Exposure to THC and Alcohol in Pregnant Rats: Maternal Outcomes and Pharmacokinetic Effects. bioRxiv, 2020.2007.2024.220103.

Cooper, S.Y., Akers, A.T., Henderson, B.J., 2020. Flavors enhance nicotine vapor self-administration in male mice. Nicotine Tob Res.

Crummy, E.A., O’Neal, T.J., Baskin, B.M., Ferguson, S.M., 2020. One Is Not Enough: Understanding and Modeling Polysubstance Use. Front Neurosci 14, 569.

Dugas, E.N., Sylvestre, M.P., O’Loughlin, J., 2020. Type of e-liquid vaped, poly-nicotine use and nicotine dependence symptoms in young adult e-cigarette users: a descriptive study. BMC Public Health 20(1), 922.

Dunbar, M.S., Davis, J.P., Tucker, J.S., Seelam, R., Shih, R.A., D’Amico, E.J., 2020. Developmental Trajectories of Tobacco/Nicotine and Cannabis Use and Patterns of Product Co-use in Young Adulthood. Tob Use Insights 13, 1179173X20949271.

Freels, T.G., Baxter-Potter, L.N., Lugo, J.M., Glodosky, N.C., Wright, H.R., Baglot, S.L., Petrie, G.N., Yu, Z., Clowers, B.H., Cuttler, C., Fuchs, R.A., Hill, M.N., McLaughlin, R.J., 2020. Vaporized Cannabis Extracts Have Reinforcing Properties and Support Conditioned Drug-Seeking Behavior in Rats. J Neurosci 40(9), 1897–1908.

Frie, J.A., Underhill, J., Zhao, B., de Guglielmo, G., Tyndale, R.F., Khokhar, J.Y., 2020. OpenVape: an Open-Source E-Cigarette Vapour Exposure Device for Rodents. eNeuro.

Gerak, L.R., Weed, P.F., Maguire, D.R., France, C.P., 2019. Effects of the synthetic cannabinoid receptor agonist JWH-018 on abuse-related effects of opioids in rhesus monkeys. Drug Alcohol Depend 202, 33–38.

Gorfinkel, L.R., Stohl, M., Greenstein, E., Aharonovich, E., Olfson, M., Hasin, D., 2020. Is Cannabis being used as a substitute for non-medical opioids by adults with problem substance use in the United States? A within-person analysis. Addiction.

Gutierrez, A., Creehan, K.M., Taffe, M.A., 2020a. A vapor exposure method for delivering heroin alters nociception, body temperature and spontaneous activity in female and male rats. Journal of neuroscience methods, 108993.

Gutierrez, A., Harvey, E.L., Creehan, K.M., Taffe, M.A., 2020b. Repeated heroin vapor inhalation during adolescence produces long-term alterations in nociception and anxiety-like behavior in male and female Wistar rats. In preparation.

Gutierrez, A., Nguyen, J.D., Creehan, K.M., Taffe, M.A., 2020c. Female rats self-administer heroin by vapor inhalation. Pharmacol Biochem Behav 199, 173061.

Javadi-Paydar, M., Creehan, K.M., Kerr, T.M., Taffe, M.A., 2019a. Vapor inhalation of cannabidiol (CBD) in rats. Pharmacol Biochem Behav 184, 172741.

Javadi-Paydar, M., Kerr, T.M., Harvey, E.L., Cole, M., Taffe, M.A., 2019b. Effects of nicotine and THC vapor inhalation administered by an electronic nicotine delivery system (ENDS) in male rats. Drug Alcohol Depend 198, 54–62.

Javadi-Paydar, M., Nguyen, J.D., Kerr, T.M., Grant, Y., Vandewater, S.A., Cole, M., Taffe, M.A., 2018. Effects of Delta9-THC and cannabidiol vapor inhalation in male and female rats. Psychopharmacology (Berl) 235(9), 2541–2557.

Justinova, Z., Tanda, G., Redhi, G.H., Goldberg, S.R., 2003. Self-administration of delta9-tetrahydrocannabinol (THC) by drug naive squirrel monkeys. Psychopharmacology (Berl) 169(2), 135–140.

Killian, A., Bonese, K., Rothberg, R.M., Wainer, B.H., Schuster, C.R., 1978. Effects of passive immunization against morphine on heroin self-administration. Pharmacol Biochem Behav 9(3), 347–352.

Kowitt, S.D., Osman, A., Meernik, C., Zarkin, G.A., Ranney, L.M., Martin, J., Heck, C., Goldstein, A.O., 2019. Vaping cannabis among adolescents: prevalence and associations with tobacco use from a cross-sectional study in the USA. Bmj Open 9(6).

Li, J.X., McMahon, L.R., Gerak, L.R., Becker, G.L., France, C.P., 2008. Interactions between Delta(9)-tetrahydrocannabinol and mu opioid receptor agonists in rhesus monkeys: discrimination and antinociception. Psychopharmacology (Berl) 199(2), 199–208.

Lichtman, A.H., Martin, B.R., 1990. Spinal action of cannabinoid-induced antinociception. NIDA Res Monogr 105, 422–424.

Maguire, D.R., France, C.P., 2018. Reinforcing effects of opioid/cannabinoid mixtures in rhesus monkeys responding under a food/drug choice procedure. Psychopharmacology (Berl) 235(8), 2357–2365.

Maguire, D.R., Yang, W., France, C.P., 2013. Interactions between mu-opioid receptor agonists and cannabinoid receptor agonists in rhesus monkeys: antinociception, drug discrimination, and drug self-administration. J Pharmacol Exp Ther 345(3), 354–362.

Manning, B.H., Merin, N.M., Meng, I.D., Amaral, D.G., 2001. Reduction in opioid- and cannabinoid-induced antinociception in rhesus monkeys after bilateral lesions of the amygdaloid complex. J Neurosci 21(20), 8238–8246.

Miliano, C., Scott, E.R., Murdaugh, L.B., Gnatowski, E.R., Faunce, C.L., Anderson, M.S., Reyes, M.M., Gregus, A.M., Buczynski, M.W., 2020. Modeling drug exposure in rodents using e-cigarettes and other electronic nicotine delivery systems. Journal of neuroscience methods 330, 108458.

Miller, M.L., Creehan, K.M., Angrish, D., Barlow, D.J., Houseknecht, K.L., Dickerson, T.J., Taffe, M.A., 2013. Changes in ambient temperature differentially alter the thermoregulatory, cardiac and locomotor stimulant effects of 4-methylmethcathinone (mephedrone). Drug Alcohol Depend 127(1-3), 248–253.

Montanari, C., Kelley, L.K., Kerr, T.M., Cole, M., Gilpin, N.W., 2020. Nicotine e-cigarette vapor inhalation effects on nicotine & cotinine plasma levels and somatic withdrawal signs in adult male Wistar rats. Psychopharmacology (Berl) 237(3), 613–625.

Moore, C.F., Davis, C.M., Harvey, E.L., Taffe, M.A., Weerts, E.M., 2020a. Antinociceptive, hypothermic, and appetitive effects of vaped and injected Δ9-tetrahydrocannabinol (THC) in rats: exposure and dose-effect comparisons by strain and sex. bioRxiv, 2020.2010.2006.327312.

Moore, C.F., Stiltner, J.W., Davis, C.M., Weerts, E.M., 2020b. Translational models of cannabinoid vapor exposure in laboratory animals. Behavioural Pharmacology Publish Ahead of Print.

Moussawi, K., Ortiz, M.M., Gantz, S.C., Tunstall, B.J., Marchette, R.C.N., Bonci, A., Koob, G.F., Vendruscolo, L.F., 2020. Fentanyl vapor self-administration model in mice to study opioid addiction. Science Advances 6(32), eabc0413.

Musshoff, F., Trafkowski, J., Lichtermann, D., Madea, B., 2010. Comparison of urine results concerning co-consumption of illicit heroin and other drugs in heroin and methadone maintenance programs. Int J Legal Med 124(5), 499–503.

Nguyen, J.D., Aarde, S.M., Cole, M., Vandewater, S.A., Grant, Y., Taffe, M.A., 2016a. Locomotor Stimulant and Rewarding Effects of Inhaling Methamphetamine, MDPV, and Mephedrone via Electronic Cigarette-Type Technology. Neuropsychopharmacology 41(11), 2759–2771.

Nguyen, J.D., Aarde, S.M., Vandewater, S.A., Grant, Y., Stouffer, D.G., Parsons, L.H., Cole, M., Taffe, M.A., 2016b. Inhaled delivery of Delta(9)-tetrahydrocannabinol (THC) to rats by e-cigarette vapor technology. Neuropharmacology 109, 112–120.

Nguyen, J.D., Bremer, P.T., Hwang, C.S., Vandewater, S.A., Collins, K.C., Creehan, K.M., Janda, K.D., Taffe, M.A., 2017. Effective active vaccination against methamphetamine in female rats. Drug Alcohol Depend 175, 179–186.

Nguyen, J.D., Creehan, K.M., Grant, Y., Vandewater, S.A., Kerr, T.M., Taffe, M.A., 2020. Explication of CB1 receptor contributions to the hypothermic effects of Delta(9)-tetrahydrocannabinol (THC) when delivered by vapor inhalation or parenteral injection in rats. Drug Alcohol Depend 214, 108166.

Nguyen, J.D., Grant, Y., Creehan, K.M., Hwang, C.S., Vandewater, S.A., Janda, K.D., Cole, M., Taffe, M.A., 2019. Delta(9)-tetrahydrocannabinol attenuates oxycodone self-administration under extended access conditions. Neuropharmacology 151, 127–135.

Nguyen, J.D., Grant, Y., Kerr, T.M., Gutierrez, A., Cole, M., Taffe, M.A., 2018a. Tolerance to hypothermic and antinoceptive effects of 9-tetrahydrocannabinol (THC) vapor inhalation in rats. Pharmacol Biochem Behav 172, 33–38.

Nguyen, J.D., Hwang, C.S., Grant, Y., Janda, K.D., Taffe, M.A., 2018b. Prophylactic vaccination protects against the development of oxycodone self-administration. Neuropharmacology 138, 292–303.

Nicksic, N.E., Do, E.K., Barnes, A.J., 2020. Cannabis legalization, tobacco prevention policies, and Cannabis use in E-cigarettes among youth. Drug Alcohol Depend 206, 107730.

Olthuis, J.V., Darredeau, C., Barrett, S.P., 2013. Substance use initiation: the role of simultaneous polysubstance use. Drug Alcohol Rev 32(1), 67–71.

Panlilio, L.V., Justinova, Z., Goldberg, S.R., 2010. Animal models of cannabinoid reward. British Journal of Pharmacology 160(3), 499–510.

Parsons, L.H., Hurd, Y.L., 2015. Endocannabinoid signalling in reward and addiction. Nature reviews. Neuroscience 16(10), 579–594.

Pearson, J.L., Villanti, A.C., 2020. It Is Past Time to Consider Cannabis in Vaping Research. Nicotine Tob Res 22(5), 597–598.

Peckham, E.M., Traynor, J.R., 2006. Comparison of the antinociceptive response to morphine and morphine-like compounds in male and female Sprague-Dawley rats. J Pharmacol Exp Ther 316(3), 1195–1201.

Ponzoni, L., Moretti, M., Sala, M., Fasoli, F., Mucchietto, V., Lucini, V., Cannazza, G., Gallesi, G., Castellana, C.N., Clementi, F., Zoli, M., Gotti, C., Braida, D., 2015. Different physiological and behavioural effects of e-cigarette vapour and cigarette smoke in mice. Eur Neuropsychopharmacol 25(10), 1775–1786.

Ramadan, M.M., Banta, J.E., Bahjri, K., Montgomery, S.B., 2020. Marijuana users are likely to report opioid misuse among adults over 50 years in representative sample of the United States (2002-2014). J Addict Dis, 1–8.

Reboussin, B.A., Rabinowitz, J.A., Thrul, J., Maher, B., Green, K.M., Ialongo, N.S., 2020. Trajectories of cannabis use and risk for opioid misuse in a young adult urban cohort. Drug Alcohol Depend 215, 108182.

Taffe, M.A., Creehan, K.M., Vandewater, S.A., 2015. Cannabidiol fails to reverse hypothermia or locomotor suppression induced by Delta(9) - tetrahydrocannabinol in Sprague-Dawley rats. Br J Pharmacol 172(7), 1783–1791.

Taffe, M.A., Creehan, K.M., Vandewater, S.A., Kerr, T.M., Cole, M., 2020. Effects of Delta(9)-tetrahydrocannabinol (THC) vapor inhalation in Sprague-Dawley and Wistar rats. Exp Clin Psychopharmacol.

Thrul, J., Rabinowitz, J.A., Reboussin, B.A., Maher, B.S., Ialongo, N.S., 2020. Adolescent cannabis and tobacco use are associated with opioid use in young adulthood-12-year longitudinal study in an urban cohort. Addiction.

Vendruscolo, J.C.M., Tunstall, B.J., Carmack, S.A., Schmeichel, B.E., Lowery-Gionta, E.G., Cole, M., George, O., Vandewater, S.A., Taffe, M.A., Koob, G.F., Vendruscolo, L.F., 2018. Compulsive-Like Sufentanil Vapor Self-Administration in Rats. Neuropsychopharmacology 43(4), 801–809.

Wakley, A.A., Craft, R.M., 2011. THC-methadone and THC-naltrexone interactions on discrimination, antinociception, and locomotion in rats. Behav Pharmacol 22(5-6), 489–497.

Wright, M.J., Jr., Angrish, D., Aarde, S.M., Barlow, D.J., Buczynski, M.W., Creehan, K.M., Vandewater, S.A., Parsons, L.H., Houseknecht, K.L., Dickerson, T.J., Taffe, M.A., 2012. Effect of ambient temperature on the thermoregulatory and locomotor stimulant effects of 4-methylmethcathinone in Wistar and Sprague-Dawley rats. PLoS One 7(8), e44652.

